# The Role of SARS-CoV-2 Nucleocapsid Protein Persistence in Inducing Chronic Type I Interferon and Mitochondrial Dysfunction

**DOI:** 10.1101/2025.06.02.657400

**Authors:** Andreas Koenig, Mohammad Quasem, Benjamin Zink, Doniyorbek Abdulla-Zoda, Priyanka Karunanidhi, Karolina Sienko, Christian Geier, Venkatesh Jayaraman, Rabia Malik, Kanza Ahmed, Paul F. Shanley, Iwona A. Koenig

## Abstract

The potential mechanisms that link SARS-CoV-2 infection to post-acute sequelae of SARS-CoV-2 symptoms and ultimately transition to new onset of autoimmune disease remain poorly understood. Here, we report the consequences of SARS-CoV-2 nucleocapsid (N) protein persistence in the absence of detectable SARS-CoV-2 replication in otherwise healthy individuals, a set of systemic lupus erythematosus (SLE) patients, and a case of a 60-year-old man who developed a new onset of lupus nephritis seven months following mild COVID-19 disease. We have identified that N protein persistence in peripheral blood mononuclear cells (PBMC) did not correlate with detectable SARS-CoV-2 RNA in blood but is associated with significantly increased secretion of type I interferons (IFN) and presence of tubuloreticular structures in peripheral leukocytes. We further demonstrate that the N protein colocalizes in the mitochondrial fraction of PBMCs from individuals positive for N protein alongside mitochondrial antiviral signaling protein (MAVS). In vitro expression of the N protein in a monocytic cell line showed that the N protein itself was directly capable of interacting with MAVS in the absence of viral RNA, and this interaction was enhanced if the cell was exposed to oxidative stress. We show that the type I IFN signature in the presence of N protein expression was MAVS, but not STING signaling pathway-dependent. Our findings suggest a mechanism for the onset and promotion of type I IFN signature after COVID-19; in our model, the SARS-CoV-2 N protein can independently trigger sustained type I IFN production via direct activation of MAVS and its spontaneous oligomerization. This long-lasting type I IFN generation may create a chronic inflammatory milieu, favoring autoimmunity with SLE-like symptoms in susceptible individuals.

**One Sentence Summary:** The persistence of SARS-CoV-2 nucleocapsid in the absence of viral replication is associated with an induction of mitochondria-mediated inflammatory milieu, which can favor autoimmunity in susceptible individuals.

## INTRODUCTION

While most individuals fully recover from COVID-19 and do not show any late-stage complications arising from the infection, a notable minority of about 6 out of 100 individuals, including those who initially exhibited mild symptoms and were vaccinated, may develop post-acute infection syndrome (PAIS), commonly known as long COVID [1–5]. Despite ongoing vaccinations and a generally milder presentation of acute SARS-CoV-2 infections during the later waves caused by different virus variants, the overall burden of long COVID is not decreasing; instead, it is expected to surpass the burden of other chronic diseases [6, 7]. The potential mechanisms that lead to or predispose individuals to long COVID, as well as the subsequent diagnosis of potentially causatively connected autoimmune disorders, remain poorly understood.

One of the current hypothesized mechanisms contributing to long COVID include viral persistence. Indeed, traces of SARS-CoV-2 have frequently been detected – even months after the initial infection - in organs distant from the initial pulmonary site of infection [8]. Interestingly, viral persistence, especially in circulating blood cells of long COVID patients, is often detected not at the level of infectious virus or viral RNA but rather at the level of viral proteins [9]. Specifically, the viral spike (S) and nucleocapsid (N) proteins have been found either as soluble proteins in plasma or serum or associated with myeloid cells [10–12]. However, the consequences of these long-lasting viral protein deposits on immune or epithelial cells’ function are poorly understood. Other proposed mechanisms contributing to long COVID include the reactivation of other dormant viruses and inflammation-induced chronic tissue dysfunction and damage [13]. There is growing evidence that the inflammation-mediated tissue dysfunction in individuals with long COVID can partly originate from myeloid cells carrying viral antigens that become proinflammatory [14].

Whether and how there is a causative connection and progression from long COVID to autoimmune and autoinflammatory connective tissue disorders is poorly understood. Recent retrospective studies suggest an increased prevalence of inflammatory arthritis, systemic lupus erythematosus (SLE), rheumatoid arthritis (RA), and systemic sclerosis in the three years following the onset of the COVID-19 pandemic, compared to the three years preceding the pandemic [15–17]. The increased incidence of defined autoimmune disorders is further correlated with the increased prevalence of new onset of autoantibodies being detected up to 16 months post-acute infection [18]. However, the factors determining whether a patient with long COVID-19 will recover or progress to chronic autoimmune diseases are unknown.

One of the key antibodies that seems to define both acute SARS-CoV-2 infection and long COVID-19 are autoantibodies against type I interferon (IFN), especially IFN-α [19–21]. This suggests an increased level of the IFN-α cytokine circulating either before the acute SARS-CoV-2 infection or post-infection, when it drives autoimmunity [22]. Interestingly, it is also well described that type I IFN are key cytokines linked to the development and progression of SLE, and it is associated with poorer outcomes [23, 24].

The relationship between SARS-CoV-2 and IFN-I is complex. The virus can subvert various stages of type I IFN production, and among those who develop severe COVID-19, there is a greater likelihood of the host carrying mutations in genes related to type I IFN production or producing inhibitory autoantibodies against type I IFN [20, 25, 26]. On the contrary, other studies have suggested that type I IFN may be involved in the development of long COVID, and such a finding could also be highly relevant for promoting the pathogenesis of SLE or other autoimmune diseases [27]. Whether the persistence of S and N proteins in circulation is associated with increased IFN-α is unknown.

Here, we present an analysis of N and S protein presence in peripheral blood monocytic cells isolated from a cohort of healthy individuals before the pandemic, within three months of acute SARS-COV-2 infection, 12 months post-initial infection, and two vaccinations. We compare the frequency of S and N protein presence in healthy individuals and the presence of viral antigens in a cohort of SLE patients. Importantly, we present our findings from the case of a new-onset SLE patient several months after an episode of mild COVID-19 in an unlikely individual, a twice-vaccinated 60-year-old man who had never been seriously ill. We demonstrate that de novo diagnosed SLE patients had increased levels of IFN-α in circulation, which correlated with N protein interaction with MAVS and its spontaneous oligomerization compared to healthy controls. We have identified that in a de novo post-COVID SLE case, evidence of type I IFN signature could also be observed in PBMCs, as detected by SLE-characteristic tubuloreticular inclusions (TRIs). In a model monocytic cell line, we finally demonstrate that the N-protein alone, in the absence of viral RNA, can trigger MAVS-dependent type I IFN production, which is further augmented by increased oxidative stress.

## RESULTS

### N protein persistence and type I IFN signature in newly diagnosed SLE patient

We recently documented a case of “long COVID” that progressed into severe de novo lupus nephritis in a previously healthy male, who had not been hospitalized or consulted a physician for over 20 years (ref). Among the many factors that led to the diagnosis of lupus in the index patient was a significant type I IFN signature detected in the kidney biopsy in the form of TRIs and in the serum as increased circulatory type IFN-α and IFN-β, when compared to healthy controls (ref). This raised the question of whether persistent SARS-CoV-2 could be responsible for long COVID in the index patient and its type I IFN signature.

We investigated the presence of SARS-CoV-2 on the N and S proteins and RNA level in the serum and PBMCs of the de novo diagnosed SLE patient and matched healthy controls (HCs). The HCs had the same vaccination status as the index SLE patient and experienced mild COVID-19 within a similar timeframe. The immunoblotting revealed that the SLE patient had significantly elevated levels of N but not S protein in the serum and the PBMCs during SLE diagnosis compared to two HCs (Fig. 1A, B). The SARS-CoV-2 RNA could not be detected in the SLE patient or HCs (Fig. S1). The same analysis conducted one and a half years later, during which the SLE patient experienced a subsequent onset of acute COVID-19 and reactivation of the varicella zoster virus, followed by a month-long treatment regimen with antiviral medications, revealed a significant reduction in the N protein level (Fig. 1A, B, C). The identical two HCs in the second timepoint study, although they also experienced another bout of COVID-19, displayed a total lack of N protein (Fig. 1A, B).

**Fig. 1.**
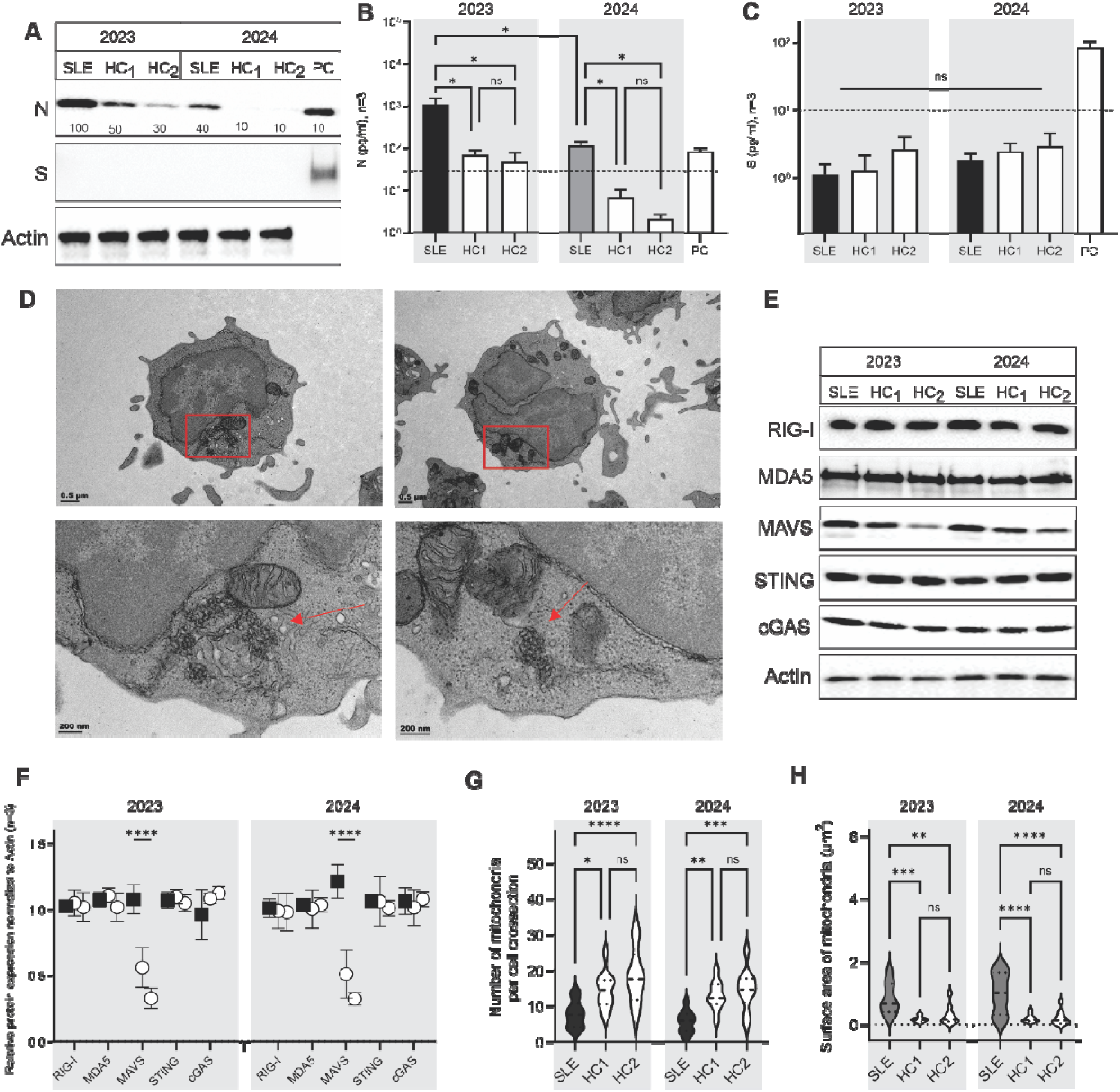
The persistence of N protein in peripheral blood monocytes of de novo diagnosed SLE patients and its association with the type I IFN signature. (A) Original immunoblots and (B, C) the corresponding relative quantification of SARS-CoV-2 nucleocapsid (N) and spike (S) protein analyzed in an equal number of peripheral blood monocytes isolated from the de novo diagnoses SLE patient and matching healthy controls (HC) within 9 and 15 months following the initially documented acute infection. (D) Transmission electron microscopy images of PBMC from SLE patient showing tubuloreticular structures (TRI), highlighted by red arrows, as a signature of type I IFN. (E, F) Immunoblotting analysis at 9- and 15-month post-infection of cytoplasmic RNA and DNA sensors and their downstream receptors in whole cell lysates of PBMCs. The expression of each protein for the first replicate of the SLE patient was assigned the value of one, and the relative expression for other SLE replicates and HCs was calculated, respectively. (G, H) Quantitative analyses of the number of mitochondria (G) and total mitochondrial area (H) per square micron. Images of 150 single cells, across random grids prepared from a PBMC cell pellet of SLE and matching HCs, were imaged by TEM. The statistical difference between the groups was analyzed by ordinary two-way ANOVA and the Tukey’s multiple comparisons test.

Next, we investigated whether the previously published pronounced type I IFN signature observed in the kidney of the index SLE patient in the form of TRIs could also be observed in the PBMCs. We have determined that approximately 6% of single cells in the de novo diagnosed SLE patient were positive for TRIs (Fig. 1D). We could not detect TRIs in cells of the corresponding healthy controls (Fig. 1S). Interestingly, 90% of cells positive for the TRIs showed a clear association of these structures with mitochondria (Fig. 1D).

Two innate immune pathways leading to a type I IFN signature are associated with the mitochondria and endoplasmic reticulum in the cells’ cytoplasm. Our analysis suggests that there are no significant differences in the expression of cytoplasmic helicases, which can recognize the self-RNA or viral RNA of SARS-CoV-2, including the retinoic acid-inducible gene I (RIG-I) and melanoma differentiation-associated gene 5 (MDA5) (Fig. 1E). Similarly, we did not identify significant differences in the expression of Stimulator of Interferon Genes (STING), cyclic GMP-AMP synthase (cGAS) (Fig. 1E). We also did not identify changes in overall expression of toll-like receptors (TLRs) −2, −3, −4, −7, and −8; the downstream signaling adaptor molecules, myeloid differentiation factor 88 (MyD88) and Toll/IL-1 receptor domain-containing adaptor inducing IFN-β (TRIF; also known as TICAM-1) (Fig. S1). By contrast, the expression of the common downstream adaptor protein of the RIG-I helicases, MAVS, was significantly increased in the PBMCs of the de novo diagnosed SLE patient compared to HCs, which did not differ significantly from each other (Fig. 1E).

### Correlation of N and S protein presence with IFN-I signature in SLE patients

The effect of the persistence of SARS-CoV-2 protein antigens on the type I IFN signature in autoimmune patients is unknown. We have investigated the presence of N or S protein in the PBMC of SLE patients routinely followed at the Upstate Medical University Rheumatology Clinic. The cohort of SLE patients consisted of samples collected before the SARS-CoV-2 pandemic (n = 7), which were previously characterized for type I IFN signature and MAVS activation [28]. The next set of samples (n = 7) was collected during the COVID-19 pandemic, between 2020 and 2022. The third set of samples (n = 7) was collected between 2023 and 2025. Each SLE subject was matched with a corresponding healthy control individual of the same age and sex.

We did not detect the presence of N and S proteins in the PBMCs of SLE patients and their matched healthy controls, whose blood was collected in 2016 (Fig. 2A-D). In the pandemic cohort of SLE patients and healthy controls, we identified only one SLE patient and one healthy individual positive for N. Still, neither was positive for S, as determined by immunoblotting and ELISA (Fig. 2A-D). In the late pandemic cohort, we identified the N protein in three SLE patients and three healthy controls (Fig. 2A-C). In this cohort, we also identified S protein in the PBMC of one SLE patient and two healthy controls. Within the three positive S samples, only one SLE patient and one healthy control were simultaneously positive for N (Fig. 2A-D). The detected S was 100 kDa for all cases, corresponding to a cleaved form, not a full-length protein (Fig. 2C, D).

**Fig. 2.**
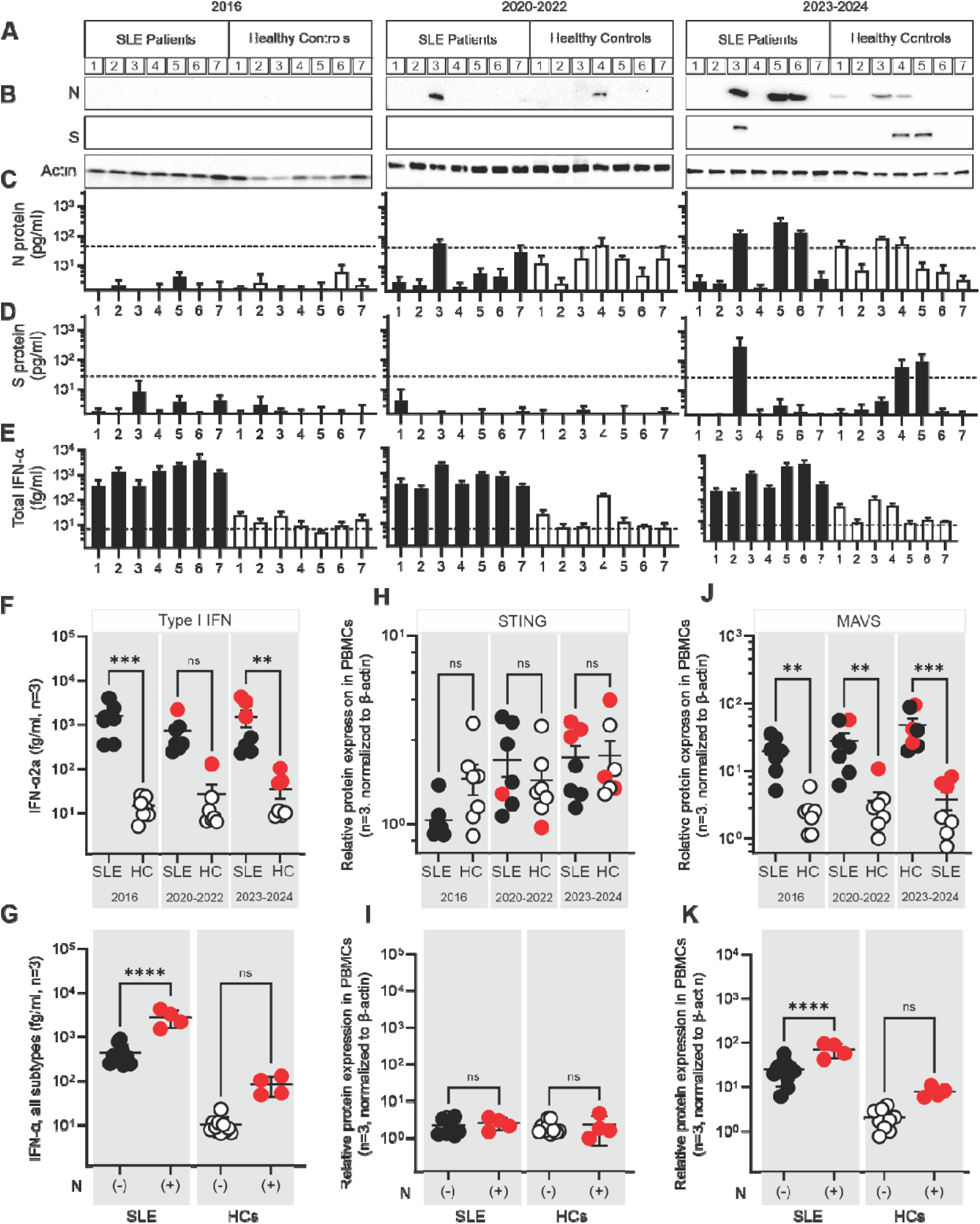
Correlation between SARS-CoV-2 N and S protein presence, type I IFN signature, and STING/MAVS expression in PBMCs from established SLE patients and matched healthy controls (HCs). (A) Blood samples were collected from SLE patients and age- and sex-matched HCs during three time periods: pre-pandemic (2016), pandemic (2020–2022), and post-pandemic (2023–2024). (B) The presence of viral nucleocapsid (N) and spike (S) proteins in PBMCs was assessed by immunoblotting, and (C, D) quantitatively measured by ELISA, with bars showing mean ± SD from three technical replicates; dotted lines indicate the lower detection limit. (E) IFN-α2 levels were measured using the S-PLEX Human IFN-α2a kit (lower detection limit 4.9 fg/ml), with three technical replicates averaged and plotted as mean ± SD. (F, G) Total type I IFN levels were stratified by N protein status, with N-positive samples highlighted as red-filled circles. (H–K) STING and MAVS protein expression in PBMC lysates was normalized relative to the first technical replicate (set to one), with mean ± SD plotted for the remaining replicates and samples. Statistical differences were assessed using two-way ANOVA with Tukey’s multiple comparisons test; *P < 0.05, **P < 0.005, ns = not significant.

Next, the correlation between type I IFN signature and the presence or absence of N and S proteins in the PBMCs of SLE patients and matched healthy controls was assessed. We have identified that N, but not S, is associated with at least a threefold increase in the total IFN-α level in plasma. The N protein-dependent increase in IFN-α was observed in SLE patients and HCs. The N-positive samples clustered in value in SLE patients and HCs, respectively (Fig. 2E-G). The level of type I IFN in SLE patients positive for N protein was significantly higher when compared to SLE patients negative for N protein (Fig. 2E-G).

In the index SLE patient, the IFN-α signature and N protein positivity correlated with increased expression of MAVS but not STING protein. Therefore, we examined whether the levels of these cytokines change significantly in the presence of N protein and whether the change is different between SLE patients and HCs. The analysis indicates that the expression of STING was randomly distributed between the individuals belonging to the SLE or HC group, regardless of N or S protein presence (Fig. 2H-I). Furthermore, no statistical difference was observed between the groups when all subjects, SLE and HCs, were stratified for N presence (Fig. 2G). In contrast, MAVS expression in the PBMCs of SLE patients and healthy controls was significantly different (Fig. 2J-K). MAVS expression in the PBMCs of SLE and HCs, which were positive for N protein, was significantly increased compared and paralleling the increased IFN-α levels in these samples (Fig. 2G and K).

### Presence of N and S proteins in PBMCs of healthy individuals and their correlation with IFN-**α** cytokine secretion and expression of STING and MAVS

To identify whether the persistence of N and S protein depends on the presence of autoimmune component we have capitalized upon an existing collection of samples obtained by the SUNY Upstate University Global Health Institute, which were previously used to examine the development of SARS-CoV-2-specific T and B cell immunity and establish assays to simultaneously analyze antigen-specific B and T cells after SARS-CoV-2 infection and vaccination [29–31]. This cohort comprised three groups: the first pre-pandemic group consisted of 10 females and 10 males, with an average age of 25, whose blood samples were collected in 2016 (Fig. 3A). The second group consisted of 20 females and 20 males with an average age of 43, who had mild, PCR-confirmed SARS-CoV-2 infection. They had a peripheral blood draw in 2020, on average, 50 days following symptom resolution, as confirmed by a negative PCR test (Fig. 3A). The third cohort comprised 20 female and 13 male individuals with an average age of 45, who were followed for at least 9 months after a negative SARS-CoV-2 PCR test and had received two mRNA vaccines during this period. The blood draw for this group was conducted in 2021, at least 12 weeks after the second vaccination to ensure there was no vaccine-driven inflammation (Fig. 3A).

**Figure 3.**
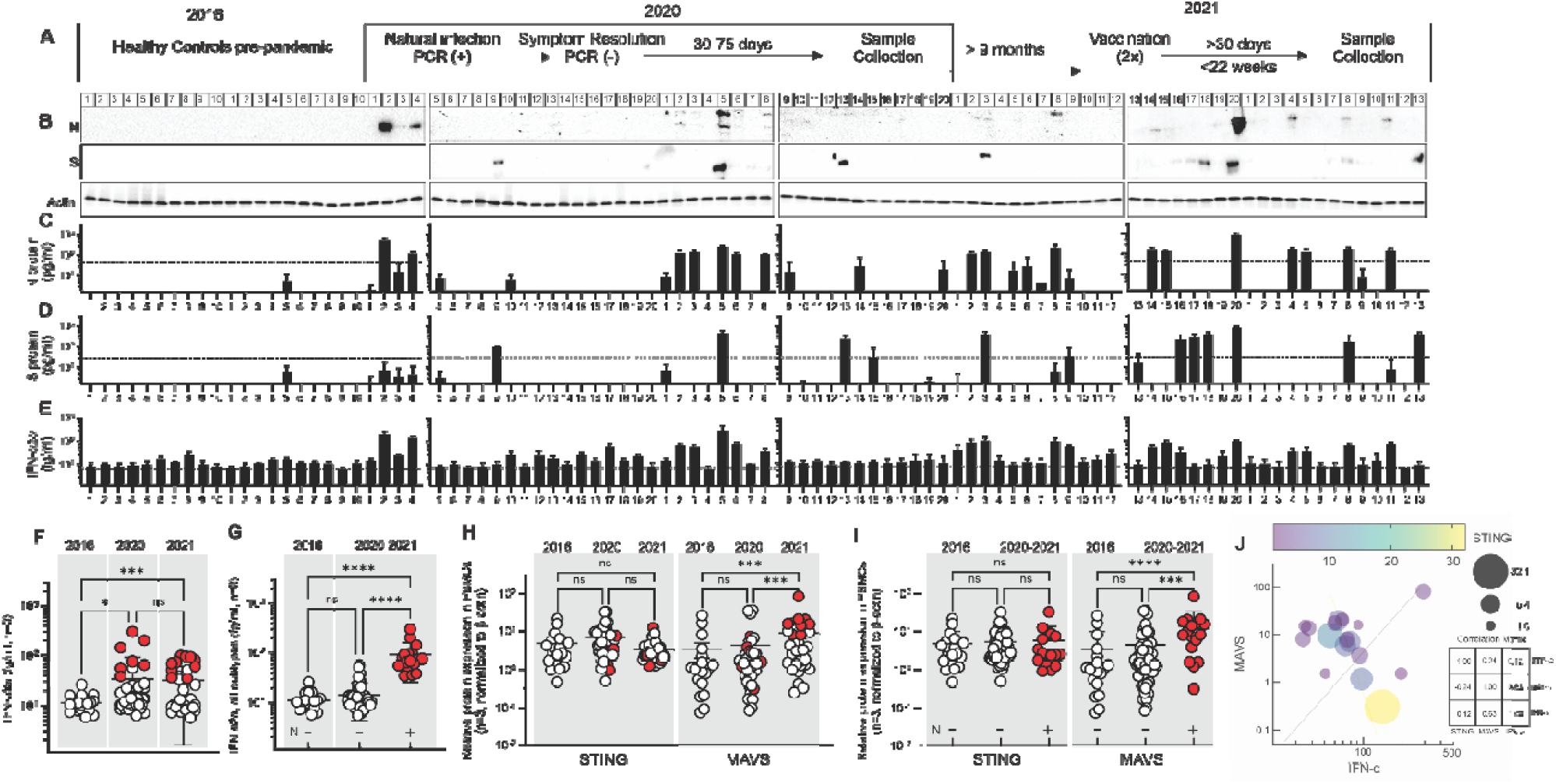
Longitudinal analysis of SARS-CoV-2 N and S protein detection, type I IFN levels, and STING/MAVS expression in PBMCs from healthy individuals pre- and post-pandemic. (A) Blood samples were collected from healthy controls (HCs; n = 20) prior to the COVID-19 pandemic (2016), during natural SARS-CoV-2 infection in 2020 (n = 40), and following vaccination in 2021 (n = 33). (B) PBMC lysates were analyzed by immunoblotting to detect SARS-CoV-2 nucleocapsid (N) and spike (S) proteins, using β-actin as a loading control. (C, D) Quantitative N and S protein abundance analysis was performed using protein-specific ELISA assays; bars represent mean ± SD across technical replicates, with dotted lines indicating lower detection limits. (E) IFN-α2 levels were quantified using the S-PLEX Human IFN-α2a kit; bars show mean ± SD across three technical replicates, with dotted lines indicating the lower detection limit. (F) Comparative analysis of IFN-α2 levels across the three cohorts, with N-positive samples highlighted as red-filled circles. (G) Comparative analysis of IFN-α2 levels between the pre-pandemic cohort and combined pandemic cohorts, with N-positive samples (red-filled circles) plotted separately. (H, I) Relative expression of STING and MAVS proteins in PBMC lysates, normalized to β-actin, plotted across cohorts and stratified by N protein status. (J) Correlation matrix illustrating the relationships among IFN-α levels, MAVS expression, and STING expression, visualized as a bubble plot where color denotes correlation strength and bubble size reflects the correlation metric. Statistical comparisons were performed using two-way ANOVA with Tukey’s multiple comparisons test; *P < 0.05, **P < 0.005, ***P < 0.001, ****P < 0.0001, ns = not significant.

We have analyzed the presence of N and S in the same manner as proteins for SLE patients and HCs using immunoblotting and ELISA on an equal number of PBMC lysates after 24 hours of culture. We did not detect protein N or S in the lysates of individuals whose blood was collected in 2016 (Fig. 3A-D). In the early pandemic cohort of healthy controls, 18% of PBMC samples tested positive for N protein and 5% for S protein. When PBMCs were collected a year later, 30% of subjects showed a positive signal for N protein, while 7% were positive for S protein (Fig. 1A-D). We detected S protein with an apparent molecular weight of 100kDa corresponding to the S1 domain but not full length (Fig. S1A). Between the two cohorts, PBMCs of 17 individuals (23%) out of 73 screened were positive for N protein, and 10 (14%) were positive for S protein. Only four samples were simultaneously positive for both N and S proteins (Fig. 1A-D). To ensure that the immunoblotting results are unambiguous, additional antibodies specific to different regions of S and N were used (Fig. S1A).

Next, we established the correlation of IFN-α with the presence of SARS-CoV-2 protein antigens. Our analysis indicated that the pre-pandemic group, testing negative for N or S protein, had an average IFN-α level significantly lower than the two pandemic groups, which were not different from each other (Fig. 3E, F). When the IFN-α values for N positive samples were subtracted from the pandemic cohorts, there was no significant difference in the IFN-α level between the pre-pandemic and pandemic cohorts of HCs (Fig. 3G). The average level of IFN-α for all N-positive samples was at least an order of magnitude higher than that observed for N-negative HCs samples (Fig. 3G).

Next, we analyzed the expression of two key cytoplasmic proteins involved in inducing type I IFN secretion. STING protein expression showed no statistically significant differences across all HC cohorts, regardless of N protein stratification (Fig. 3H, I). In contrast, analysis of MAVS expression across the three cohorts indicated that the second pandemic cohort had significantly elevated MAVS expression compared to both the pre-pandemic and early pandemic cohorts, with N protein–positive values clustering in the upper quartile (Fig. 3H). However, when N-positive samples were separated from the combined pandemic cohorts, MAVS expression was not significantly different from the pre-pandemic cohort (Fig. 3I). Notably, MAVS protein expression in N-positive samples was significantly higher compared to both N-negative samples from the pre-pandemic and pandemic cohorts (Fig. 3I). Pearson correlation analysis revealed a strong correlation (r = 0.69) between MAVS and IFN-α, while STING showed a weak correlation (r = 0.12) (Fig. 3J)

### N protein presence in PBMC is associated with the expansion of DN T cells and CD163+ monocytes

The type I IFN signature in SLE patients is often associated with changes in the relative ratio of lymphocyte sub-populations [32]. We have phenotyped the PBMCs of healthy individuals in the presence or absence of N protein. Further, we profiled in the same manner the PBMCs of the de novo diagnosed SLE patient and corresponding healthy controls at the time of the diagnosis and one year later, when the levels of N protein were decreased.

The flow cytometry analysis suggests that the N protein in the circulation of healthy individuals did not significantly affect the overall numbers of CD4+ and CD8+ T cells (Fig. 4A, B). Interestingly, N protein presence in the PBMCs was associated with a significant increase in the number of double-negative (DN) T cells (Fig. 4C). A similar increase was, however, not observed for CD19+ B cells (Fig. 4D). The same analysis performed for the total number of CD14+ monocytes suggests that individuals with N persistence had a significant lower number of these cells (Fig. 4E). However, within the CD14 positive population, the number of CD16^++^ monocytes (Fig. S4) as well as CD163^+^ monocytes (Fig. 4F) were significantly increased.

**Fig. 4.**
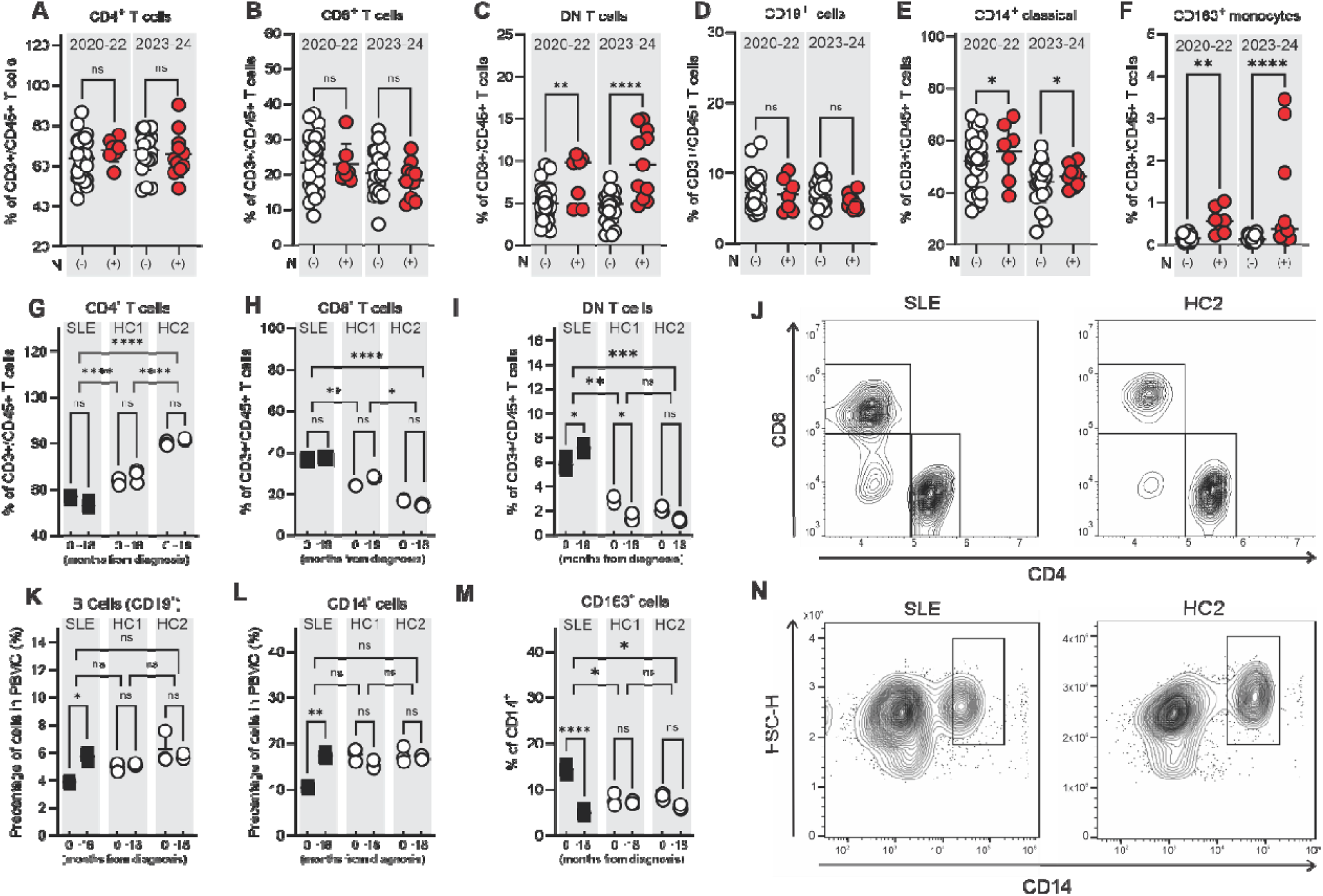
Frequency of immune cell subsets in PBMCs from SLE patients and healthy controls, stratified by nucleocapsid (N) protein presence. (A–C) Percentages of CD4, CD8, and double-negative (DN; CD4 CD8) T cells were quantified from CD3 CD45 PBMC populations in samples collected during 2020–2022 and 2023–2024, stratified by nucleocapsid (N) protein status (N (-): open circles; N(+): red-filled circles). (D–F) Percentages of CD19 B cells, CD14 classical monocytes, and CD163 monocytes, with CD163 cells subgated from the CD14 population. (G–I) Comparative analysis of CD4, CD8, and DN T cell frequencies in the de novo diagnosed SLE patient and two healthy control groups (HC1, HC2), plotted by months from diagnosis (0–18 months). (K–M) Comparative analysis of CD19 B cells, CD14 monocytes, and CD163 monocytes in the same groups over time. (J, N) Representative flow cytometry contour plots illustrating gating strategies for CD4^-^/CD8^-^ T cell subsets (J) and CD14^+^/CD163^+^ monocyte populations (N) from the index SLE patient and HC2 control. Statistical analyses were performed using ANOVA with Tukey’s multiple comparisons test; *P < 0.05, **P < 0.01, ***P < 0.001, ****P < 0.0001, ns = not significant.

The same analysis performed for the de novo diagnosed SLE patient and its matching healthy controls suggests that changes in the level of N protein in the PBMC do not affect the overall number of CD4^+^ or CD8^+^ T cells (Fig. 4G, H). However, there was a significant reduction in the overall count of CD4^+^ T cells for SLE patient compared to HC1 and HC2. (Fig. 4G). The opposite was observed for CD8^+^ T cells, which were significantly higher in number in the SLE patient than in HC1 and HC2 (Fig. 4H). DN T cells were also increased in the SLE patient compared to HCs (Fig. 4I, J). Unlike what was observed for CD4 and CD8 T cells for HC1 and HC2, which showed differences in the overall number, there was no difference between HC1 and HC2 for DN T cells (Fig. 4G-J). We have not observed a difference in the number of total B cells between the SLE and HCs (Fig. 4K).

Analysis of the monocytic compartment as defined by CD14 suggests that the SLE patient’s total number of cells expressing CD14 at the time of diagnosis was significantly lower compared to HC1 and HC2 (Fig. 4L). One year after diagnosis, the number of CD14-positive cells in the SLE patient’s PBMCs seemed to recover and reach the level observed in HC1 and HC2 (Fig. 4L). Similarly, recovery was observed for CD163+ positive cells, which were increased in the SLE patient at diagnosis and decreased one year later to the level observed in the HC1 and HC2 (Fig. 4L, M, N). This suggests that N protein presence is associated with the expansion of DN T cells and CD163 monocytes.

### N protein interacts with MAVS in the absence of viral replication in human PBMCs

To define the mechanism of how N protein can be associated with increased MAVS expression and therefore increased type I IFN in circulation, we first tested whether these two proteins interact with each other and whether this association is mediated by the interaction of MAVS with RIG-I and MDA-5 helicases and self or viral RNA. First, we performed purification of mitochondrial fraction from PBMCs of the de novo diagnosed SLE patient and matching HCs and have identified that, unlike what we had predicted, RIG-I and MDA5 were absent from the fraction (Fig. 5A). MAVS levels in the mitochondrial fraction were significantly elevated in the SLE patient compared with HCs and correlated with increased levels of phosphorylated TBK1 and IRF3 and presence of N protein (Fig. 5A). We have confirmed the specificity of this finding by detecting N protein and simultaneously phosphorylated forms of IRF3 and TBK1 only in the SLE but not in HC1 and HC2 following immunoprecipitation performed with an antibody specific to human MAVS (Fig. 5B).

**Fig. 5.**
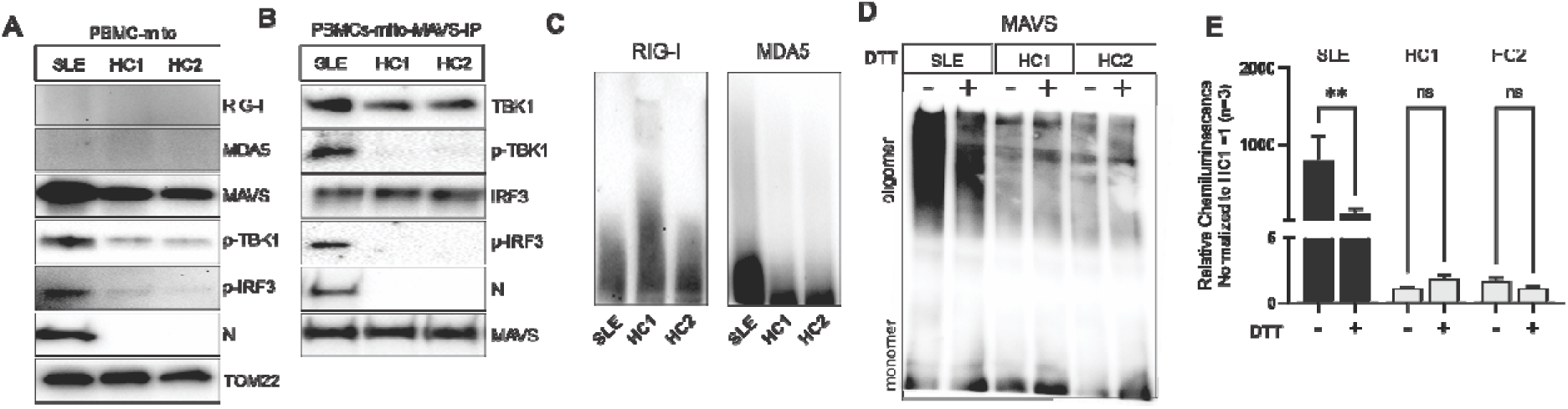
Association of SARS-CoV-2 N protein with MAVS and MAVS-pathway components in PBMCs from a de novo diagnosed SLE patient and two matched healthy controls (HCs). (A) Mitochondria were purified from PBMC lysates using a magnetically labeled antibody against translocase of the outer mitochondrial membrane 22 (TOM22). Mitochondrial fractions were probed for the cellular helicases RIG-I and MDA5, MAVS, phosphorylated downstream effectors (p-TBK1, p-IRF3), and SARS-CoV-2 N protein. TOM22 served as an internal loading control. (B) To assess the specificity of N protein association with MAVS, mitochondrial fractions were immunoprecipitated with a MAVS-specific antibody (E3) and examined for both non-activated and phosphorylated forms of TBK1 and IRF3, with MAVS serving as an internal control for immunoprecipitation efficiency. (C) High-molecular-weight oligomers of RIG-I and MDA5 were analyzed by non-denaturing gel electrophoresis and immunoblotting. (D) MAVS oligomerization was assessed using non-denaturing gel electrophoresis, probing for monomeric and oligomeric forms in the presence or absence of the reducing agent dithiothreitol (DTT). (E) Quantification of MAVS chemiluminescence signals was plotted, comparing SLE and HC samples under reducing (+DTT) and non-reducing (−DTT) conditions to evaluate differences in MAVS oligomerization. Statistical analysis was performed using two-way Anova, with Tukey multiple comparison test; **P < 0.005, ns = not significant.

RIG-I, MDA-5, and MAVS activation, whether spontaneous or induced by viral or host RNA are characterized by the formation of higher-order oligomers [33, 34]. We did not detect high molecular weight RIG-I and MDA5 oligomers in the PBMCs of SLE patients or the HCs (Fig. 5C). MAVS oligomerization, on the other hand, was readily detectable in the SLE patient but not in HCs (Fig. 5D, E). MAVS oligomerization in SLE patient was diminished when the cell extract was treated with a reducing agent (Fig. 5D, E).

### SARS-CoV-2 N protein promotes MAVS oligomerization in the absence of viral RNA

To investigate the role of the N-protein in human cells in an unbiased manner, that is, in the absence of any viral RNA and viral replication, we employed human embryonic kidney cells (HEK-293) and THP monocytic cell lines. The HEK-293 and THP cells were stably transfected with a commercially available pCDH-CMV-MCS-EF1α-Puro cloning and expression lentivector containing an inserted sequence of SARS-CoV-2 N protein. We used HEK and THP cells transfected with an empty vector or a vector carrying only GFP expression as a control. We found that N protein expression in the absence of viral RNA in HEK cells had a significant impact on the expression of MAVS, but not pattern recognition receptors RIG-I, MDA5, cGAS, or its downstream receptor STING (Fig. 6A, B). The increased expression of MAVS in cells carrying N-protein was associated with elevated secretion of IFN-α and -β (Fig. 6C, D). The secretion of IL-6 exhibited a similar pattern; however, the differences did not reach statistical significance (Fig. 6E). Furthermore, the transient transfection of HEK cells with N-protein mRNA induced a similarly detectable expression of N-protein and an associated increase of MAVS expression (Fig. S6).

**Fig. 6.**
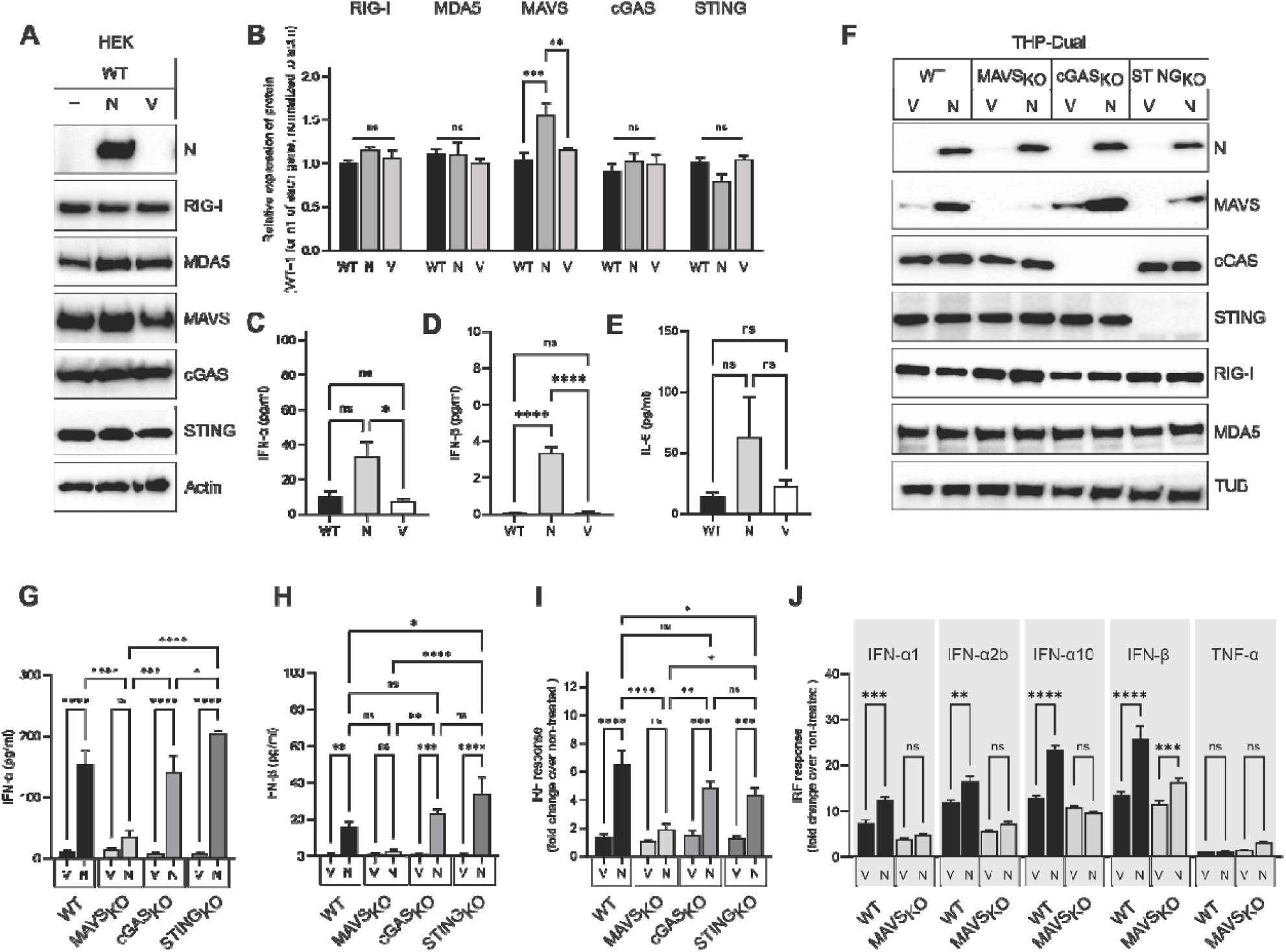
Functional Dissection of MAVS, cGAS, and STING Pathways Under SARS-CoV-2 N Protein Expression. (A) HEK293 cells were stably transfected to express SARS-CoV-2 nucleocapsid (N) protein constitutively. Lysates from N-transfected (N), non-transfected (−), and empty vector– transfected (V) cells were probed by immunoblotting for helicases RIG-I and MDA5, MAVS, and related cGAS and STING pathway components. Actin served as a loading control. (B) Relative protein expression was quantified from chemiluminescence signals normalized to actin. (C–E) Secreted cytokine levels (IFN-α, IFN-β, and IL-6) were measured in N-transfected, V-transfected, and non-transfected cells by ELISA. (F) The THP-Dual inducible reporter cell line was transfected with either N protein (N) or empty vector (V) in the background of unchanged wild-type protein expression (WT), and in the absence of MAVS (MAVS-KO), cGAS (cGAS-KO), and STING (STING-KO). THP-Dual cells were probed for expression of MAVS, cGAS, STING, RIG-I, and MDA5 relative to N protein expression; actin served as a loading control. (G, H) Secreted type I IFN levels were measured in N- and V-transfected THP-Dual cells across WT and knockout backgrounds with ELISA. (I) IRF pathway activation (fold change over non-treated) was determined under the same conditions. (J) IRF pathway activation in WT and MAVSKO THP-Dual cells was further assessed in response to stimulation with specific type I IFN-I and TNF-α. Statistical analyses were performed using 2-way ANOVA with group median ± SD; *P < 0.05, **P < 0.005, ***P < 0.0005, ****P < 0.00005, ns = not significant.

To establish whether MAVS is sufficient and necessary for N protein-mediated IFN-I signature, independently of cell type, we also used THP cells stably transfected with N protein, in the presence or absence of MAVS, STING, or cGAS expression, respectively. We confirmed the results observed in HEK cells that N-protein expression alone can lead to a significant increase in MAVS, but not RIG-I, MDA5, cGAS, or STING expression (Fig. 6F). We have further found that, with N-protein presence, only the lack of MAVS expression, but not STING or cGAS, interfered with the increased secretion of IFN-α, IFN-β, and led to loss of type I IFN signature as measured by activation of IRF responses (Fig. 4G, H, I). The same MAVS-dependence of an increase of activation of IL-6 secretion or NF-κB gene activation in cells carrying N-protein was not observed (Fig. S6). To control for the specificity of the N protein alone to be stimulatory for MAVS activation, we also transfected the cells with a lentivector carrying the viral Spike (S) protein. We identified that S protein alone, without any further stimulation, did not induce increased MAVS expression or induction of type I IFN secretion (Fig. S6).

Next, we tested whether the N protein presence in the cell also accelerated type I IFN or NF-κB gene signaling in a paracrine fashion when the cells were stimulated either with IFN-α1, IFN-α2b, IFN-α10, IFN-β, or TNF-α. We found that all four subtypes of type I IFN induced an increase in IRF response in a MAVS-dependent manner when N-protein was present (Fig. 6J). The MAVS dependence of IRF gene activation was not observed when cells were treated with TNF-α. Interestingly, in the absence of MAVS expression, N-protein had a more significant effect on the NF-κB gene expression than the WT counterpart (Fig. S6). These findings suggest that N-protein in monocyte-like cells can accelerate the type I IFN signature in both paracrine and autocrine modes.

### Reactive oxygen species accelerate SARS-CoV-2 N-protein-dependent MAVS signaling, leading to changes in mitochondrial phenotype and localization within the cell

Our group has previously identified that spontaneous, virus-independent MAVS activation leads to mitochondrial hyperpolarization and ballooning, which in turn further increases the generation of reactive oxygen species and mitochondrial dysfunction [35]. Therefore, we studied next whether exposure of the cells expressing N-protein to oxidative stress could accelerate MAVS-mediated type I IFN signaling and affect mitochondrial phenotype. We tested this by adding glucose oxidase (GOx) to culture media, which leads to continuous hydrogen peroxide production from the oxidation of β-D-glucose to D-glucono-δ-lactone [36]. We observed that the presence of N protein significantly increased MAVS / MAVS oligomerization compared to exposure to oxidative stress alone (Fig. 7A, B). In parallel, the increased oligomerization of MAVS in the presence of N-protein expression under oxidative stress correlated with a significant increase in IRF gene activation over the unstimulated cells (Fig. 7C). Furthermore, in contrast to observations in unstimulated cells, N protein expression in the presence of MAVS had a significantly higher impact on the NF-κB gene expression in the presence of oxidative stress (Fig. S7).

**Fig. 7.**
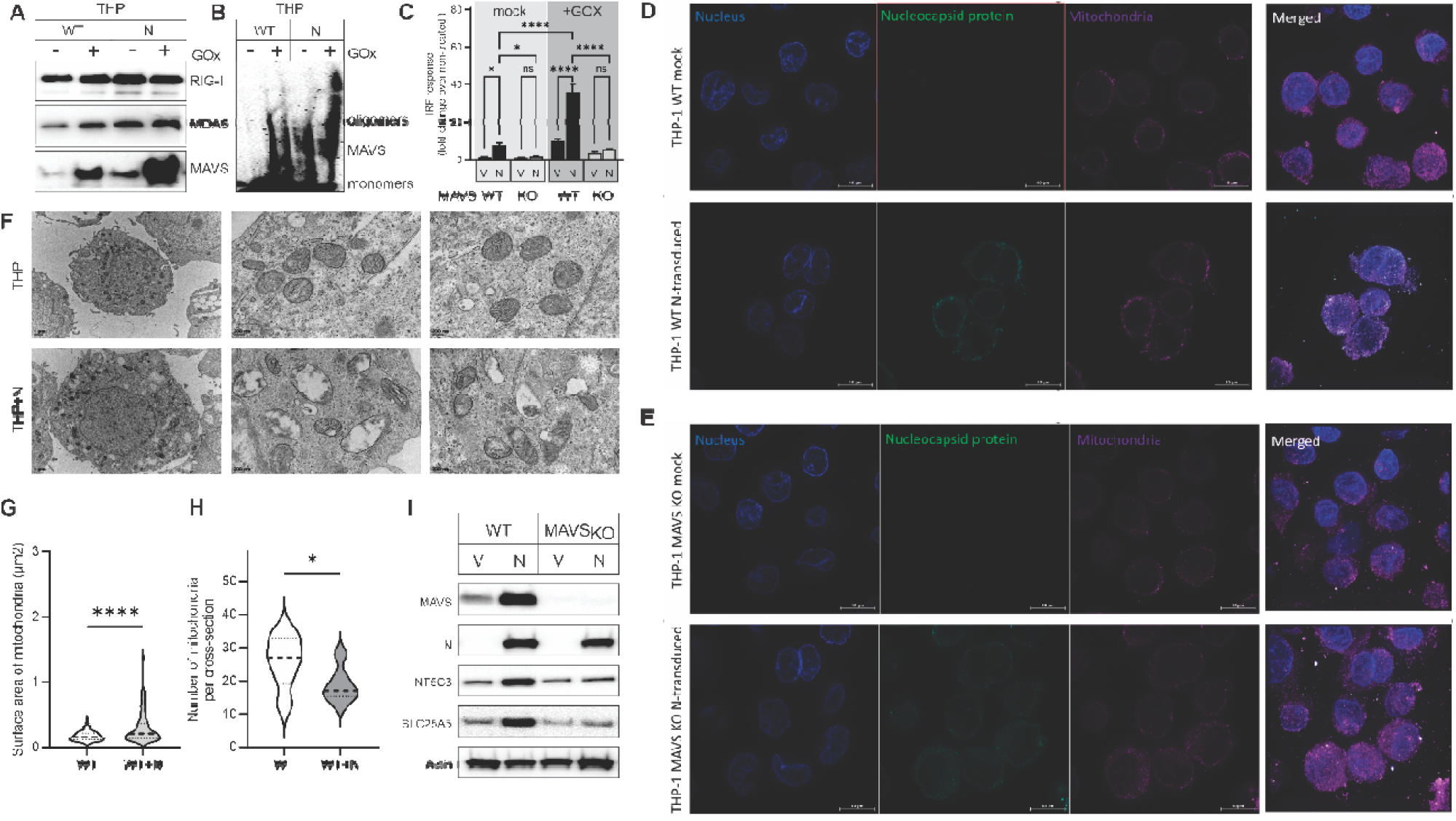
Impact of SARS-CoV-2 Nucleocapsid Protein on MAVS Oligomerization, Mitochondrial Morphology, and its Subcellular Localization depending of MAVS presence. (A) Immunoblot analysis of THP-1 wild-type (WT) cells transduced with SARS-CoV-2 nucleocapsid (N) protein or mock control (−), with or without glucose oxidase (GOx) stimulation, probed for RIG-I, MDA5, and MAVS. (B) Non-denaturing gel electrophoresis probing MAVS oligomerization states (monomer vs. oligomer) under the same conditions. (C) Quantification of IRF pathway activation (fold change over non-treated controls) across THP-1 WT and MAVS knockout (KO) cells, under N-transduced and GOx-stimulated conditions. (D, E) Confocal microscopy images of THP-1 WT and MAVS-KO cells, transduced or mock-transduced with N protein, stained for nuclei (blue), nucleocapsid protein (green), and mitochondria (purple); merged images highlight colocalization patterns. (F) Transmission electron microscopy (TEM) images showing ultrastructural mitochondrial changes in THP-1 WT cells with or without N protein expression. (G, H) Quantitative analysis of mitochondrial surface area and number per cell in WT ± N-transduced THP-1 cells. (I) Immunoblot of THP-1 WT and MAVS-KO cell lysates, with or without N transduction, probing for MAVS, N protein, NT5C3, SLC25A5, and actin as a loading control. Statistical significance was determined using two-way ANOVA; data are shown as median ± SD. *P < 0.05, **P < 0.005, ***P < 0.0005, ****P < 0.00005; ns, not significant.

N protein expression, MAVS association, and MAVS activation on mitochondria were further substantiated by confocal microscopy investigation of N-protein and mitochondrial colocalization. In THP cells expressing N-protein, the mitochondria were colocalizing with N protein in close vicinity to the outer membrane (Fig. 7D). In cells lacking MAVS expression, the N-protein was rather diffusely distributed throughout the cytoplasm of the cell and not co-localizing with mitochondria (Fig. 7D).

Lastly, we investigated whether the N-protein influenced the mitochondrial phenotype. The mitochondria in THP cells, without N protein expression, were, in most cases, distributed along the outer membrane and were spheroid in shape with well-defined cristae (Fig. 7E). In contrast, in the cells expressing N protein, the mitochondria were randomly distributed in the cytoplasm. The shape of mitochondria was more diverse ranging from spheroid to tubular, with many mitochondria lacking clearly defined cristae (Fig. 7E). Enumeration of mitochondria and their sizes suggests that the size of mitochondria in the cells expressing N protein is significantly increased with the number of mitochondria decreasing (Fig. 7F). Circling back to the PBMC of the de novo SLE patient, we observed similar phenotypical changes of mitochondrial size and number, as compared to HC1 (Fig. 7G), suggesting that N protein can induce the same changes in mitochondria as observed in the scenario of natural persistence. We have further investigated whether the expression of NT5C3, which is associated with the formation of TRIs in the SLE patient’s PBMC, was also increased in model cells expressing N protein. Indeed, we found that not only did the expression of NT5C3 increase, but also a simultaneous increase of SLC25A1 expression (Fig. 7H). This is of note as an increased expression of SLC25A4 was associated with mitochondrial dysfunction related to so-called mitochondrial clogging. Whether the interaction of MAVS and N protein can lead to mitochondrial clogging is beyond the scope of this paper but is currently under investigation.

## DISCUSSION

Despite its clinical and public health importance, persistence of SARS-CoV-2 antigens remains greatly understudied. Community-based surveillance analysis had identified that at least 0.1-0.5% of SARS-CoV-2 infections become persistent, typically with rebounding high viral loads that were documented to last for a minimum of 60 days [37]. Interestingly, SARS-CoV-2 was also shown to persist for several months as either non-replicating viral RNA or protein antigens in blood cells and plasma [38, 39]. This, along with the detectability of SARS-COV-2 in stool or urine samples of patients with persistent symptoms long after the acute infection, suggests that the virus can also be delivered to peripheral organs [10, 11, 37]. The mechanisms interconnecting SARS-CoV-2 infection, long COVID, and subsequent autoimmune disease, particularly SLE, remain largely enigmatic.

Our data suggests that while replicating SARS-CoV-2 and its viral RNA were not detectable in peripheral blood, viral nucleocapsid (N) protein was more readily detected (24% of all samples) in PBMCs, months after an initially reported infection, than S protein (12% of all samples). Although the number of SLE patients investigated was 6 times smaller than that of otherwise healthy individuals, the overall positivity for N protein was higher (33%) for SLE individuals as compared to HCs (20%) (Table 1). The same difference was not observed for persistence of S protein, which could be detected in an equal number of HCs and SLE individuals (13% and 12%).

It is not known how the persistence of SARS-CoV-2 antigen affects the function of circulating blood cells and can lead to a chronic inflammatory state in organs distal to the respiratory tract and, therefore, contribute to the onset and progression of autoimmune diseases such as SLE in predisposed individuals. Our data suggests that the persistence of protein antigens correlates with increased type I IFN in the circulation of both HC and SLE individuals, with the type I IFN being highest for SLE patients positive for N protein in circulation. The relevance of this finding correlates with recently published studies suggesting that type I IFN activity is not only critical for the acute phase of infection but also involved in the pathogenesis and symptoms of long COVID, where it is thought to promote local inflammation [43, 44].

The emphasis on type I IFN also stems from the presence of TRIs, frequently referred to in literature as “interferon footprints,” in the kidney biopsy of a patient newly diagnosed with SLE, as highlighted in our case report (ref) and the fact that we were able to observe the same structures in the PBMCs of a newly diagnosed SLE patient as well as in SLE patients and HCs positive for N protein.

TRIs are pathologic intracellular, highly structured aggregates of tubular-type structures identified by electron microscopy (EM) within the endoplasmic reticulum (ER) of endothelial cells or circulating mononuclear cells [40]. They occur naturally, either in epithelial cells of distal organs or cells of peripheral blood following various viral infections, are the hallmark of autoimmune disease diagnosis, especially lupus nephritis, or can be induced in cells exposed to type I IFN in vitro or associated with type I IFN therapy for infections or malignancy [41]. The mechanism of the TRI induction in cells is not well established, and it is not known whether the structures can be induced directly via activation of innate immune pathways in response to viral nucleic acid, or in a paracrine manner following type I IFN receptor signaling.

The TRIs were shown to contain within their structures cytosolic 5’-nucleotidase 3 (NT5C3) [40, 42]. Indeed, we found NT5C3 expression to be increased in the PBMCs of the de novo SLE patient and the model monocyte cell line expressing N protein. Why NT5C3 is a central component of TRI structures in the background of the type I IFN signature is unknown. Interestingly, NT5C3 -typically responsible for dephosphorylation of non-cyclic nucleoside 5’-monophosphates to their corresponding nucleosides, which can diffuse throughout biological membranes and are involved in a feedback loop metabolic signaling of immune cells [43–45]. Whether TRI-associated NT5C3 functions as an epigenetic regulator of cellular metabolism in the background of type I IFN signature, which is known to prevent the shift to aerobic glycolysis in inflammatory immune cells expanding in SLE and drive mitochondrial dysfunction and stress, should be further investigated [46].

Finally, our mechanistic studies aimed to clarify whether SARS-CoV-2 N-protein correlated with an excess of type I IFN production. Therefore, we examined both the association of N-protein with intracellular pathways leading to type I IFN production in the patients’ PBMCs and then in vitro by studying these pathways after specifically introducing N-protein into epithelial and monocytic cells. Previously published research suggested that the N protein targets the RIG-I pathway at multiple levels in SARS-CoV-2 infection. First, N-protein forms condensates with SARS-CoV-2 RNA, and thereby evades interaction of viral RNA with RIG-I itself; second, it directly binds to RIG-I, and third, it targets ubiquitination of MAVS and therefore its ability to aggregate, which is needed for MAVS to trigger potent type I IFN secretion [47–49]. Our results show that, in contrast to these effects during active infection, N protein augmented the expression and activation of MAVS (but not of RIG-I, MDA5, cGAS or STING) in PBMC of the SLE patient and other individuals positive for N protein expresision. We show the same MAVS activation in vitro in HEK and THP cells carrying lentiviral expression of the N-protein. We further found that N-protein and MAVS (but not RIG or MDA5) co-purified in the mitochondrial fraction of the patient’s PBMCs along with phosphorylated (activated) TBK and IRF3, the downstream components of the MAVS cascade leading to type I IFN production. We show that the same experiments performed in the absence of cGAS or STING signaling cannot abort type I IFN secretion in the presence of N protein, but that there is a loss of excess type I IFN production when MAVS is knocked out. These findings suggest that MAVS is sufficient and necessary for N-protein-mediated type I IFN production. These data correlate with confocal microscopy findings showing that the mitochondrial and N protein colocalization is enhanced by MAVS presence. Whether N protein colocalization with MAVS and it activation can lead to TRIs induction is not known and should be further investigated.

The limitations of this study are those of any studies combining cohorts of patients and healthy controls with individual case analysis. There are, very likely, factors influencing events that are either unknown or inadequately understood and cannot be isolated and controlled for. We cannot exclude the possibility that there are sources other than SARS-COV-2 to induce type I IFN signature in the de novo diagnosed SLE patient, established SLE patients, and HCs positive for N protein. There are no studies to guide thinking on what might happen with the interaction of cancer, vascular anomalies, and COVID-19 in a single patient. Further, since it is clear from our results that N-protein persists even in healthy controls, there must be additional risk factors in our patient that lead from its capacity to promote type I IFN production to uncontrolled innate immune activation and autoimmunity.

In summary, our findings suggest that the persistence of SARS-CoV-2 N-protein in peripheral blood promotes a chronic inflammatory milieu, even several months after active infection via MAVS-mediated type I IFN production that could contribute, in susceptible individuals, to the development of autoimmunity. Longitudinal studies following SLE patients and otherwise healthy individuals with or without long COVID signature but positive for protein antigens of SARS-CoV-2 could explain who, when, and why could suffer in the long term and whether therapeutics targeting mitochondria-mediated innate immunity overstimulation could bring relief.

## Notes

### Competing Interest Statement

The authors have declared no competing interest.

### Funding

- NIH: 1R01AR073282, 1R21AI128510, 1R21AR069830

